# Mechanisms behind facilitation-competition transition along rainfall gradients

**DOI:** 10.1101/2025.11.24.690218

**Authors:** Oded Hollander, Yair Mau, Niv DeMalach

## Abstract

Woody cover is rapidly changing due to mortality, shrub encroachment, and afforestation, reshaping herbaceous communities and ecosystem functioning worldwide. Often, trees and shrubs promote herb growth in dry sites but suppress it in wetter ones, as predicted by the classical Stress Gradient Hypothesis. However, explanations for the facilitation-to-competition transition remain verbal and contested, lacking a clear link to resource competition theory. Here, we present a mechanistic framework consisting of two submodels: (i) canopy shading that reduces photosynthesis and evapotranspiration, and (ii) root effects, including water uptake and increased moisture via hydraulic redistribution. We elucidate the conditions under which interactions shift from facilitation to competition. The models reproduce this reversal only when water is not the sole limiting factor at high rainfall or when woody density increases with precipitation. Moreover, the reversal can occur across any aridity gradient, including those driven by evaporative demand influenced by temperature and humidity. The two pathways leave distinct signatures: canopy shading produces a hump-shaped pattern with maximum facilitation at intermediate stress, while the root pathway predicts a shift from positive to negative interactions as water availability increases. By translating a classic idea into a quantitative framework, this model enhances ecosystem management in a changing world.

## 1 Introduction

Global climate and land-use changes are rapidly reshaping woody vegetation worldwide ^1;2^. These shifts are especially common in drylands, which cover about 40% of Earth’s land surface ^3;4^. Many water-limited systems are losing woody cover due to widespread drought and fire ^5^. Conversely, other dry regions show woody expansion, driven by shrub encroachment ^2;6^ and large-scale afforestation aimed at climate mitigation ^1;7^. In these ecosystems, woody plants (hereafter, trees) strongly shape microclimate and resource availability, thereby influencing the abundance and distribution of herbaceous plants (hereafter, herbs) that sustain forage production and biodiversity in drylands ^2;6^.

Trees are ecosystem engineers, altering the environment beyond simply consuming light and water ^8^. They can facilitate herb growth through **canopy and root mechanisms**. The canopy suppresses light availability, which reduces carbon assimilation (photosynthesis) but also lowers evapotranspiration and therefore water loss ^9^. This reduction in evapotranspiration also results from microclimatic buffering: slower wind speeds, higher humidity, and lower temperatures beneath the canopy ^10^. Tree roots not only extract water, thereby drying the soil, but can also increase soil moisture by enhancing infiltration ^11^ and redistributing water from deeper to shallower soil layers through hydraulic lift ^12;13^.

Rainfall amount often mediates the balance between these positive and negative effects. Meta-analyses find that trees typically benefit herbs at low rainfall but hinder them as rainfall increases ^14–16^. Even so, some studies report inconsistent neighbor effects along similar gradients ^17^, pointing to hidden thresholds or additional factors that alter the expected pattern.

Over the past three decades, the transition from facilitation to competition has largely been investigated through the lens of the Stress Gradient Hypothesis (SGH). This conceptual framework was proposed to explain why facilitation dominates under high abiotic stress (low rainfall), whereas competition prevails under benign conditions ^18–23^. The original explanations emphasized plant responses along water-stress gradients ^20^, but the hypothesis has since been applied to many other stress types ^24;25^. The primary argument is that tree-mediated relief of water stress dominates under low rainfall, whereas shading-induced inhibition dominates under low stress (high rainfall) ^9^. These verbal arguments were later extended using phenomenological models that embedded spatial–temporal dynamics ^26^ and biodiversity feedbacks ^27^.

Despite its prominence, the SGH has been questioned both mechanistically and in terms of predicted patterns ^9;28–31^. Some studies argue that species interactions become more strongly negative with rainfall ^14^, whereas others report a unimodal (hump-shaped) pattern in which facilitation peaks at intermediate stress and weakens under both severe aridity and benign conditions ^29^. Resolving these discrepancies calls for quantitative models that explicitly represent resource dynamics and make assumptions transparent, allowing specific processes such as shading or water uptake to be identified as drivers that generate, sustain, or limit facilitation along the gradient.

Consumer-resource theory is the leading modeling framework for mechanistic explanations of species interactions ^32–35^. Yet, it has only recently been applied to the SGH ^36;37^. These recent applications yielded insights into the role of trees in elevating resource availability during early succession ^36^ and into the joint effects of drought and grazing ^37^. However, they focused exclusively on root mechanisms and did not consider canopy shading, the mechanism emphasized in the classical conceptual hypothesis ^20^. Crucially, and contrary to the SGH’s prediction of a facilitation-to-competition shift, these consumer-resource models produced an interaction sign that remained constant (either positive or negative) along the rainfall gradient, rather than a transition from facilitation to competition.

Here, we develop a minimal consumer–resource framework that generates the classic shift from facilitation to competition as rainfall (resource supply rate) increases. The model clarifies the conditions under which this transition emerges for both canopy and root mechanisms (Fig. 1) and reconciles contrasting predictions regarding how facilitation strength varies along precipitation gradients.

**Figure 1.**
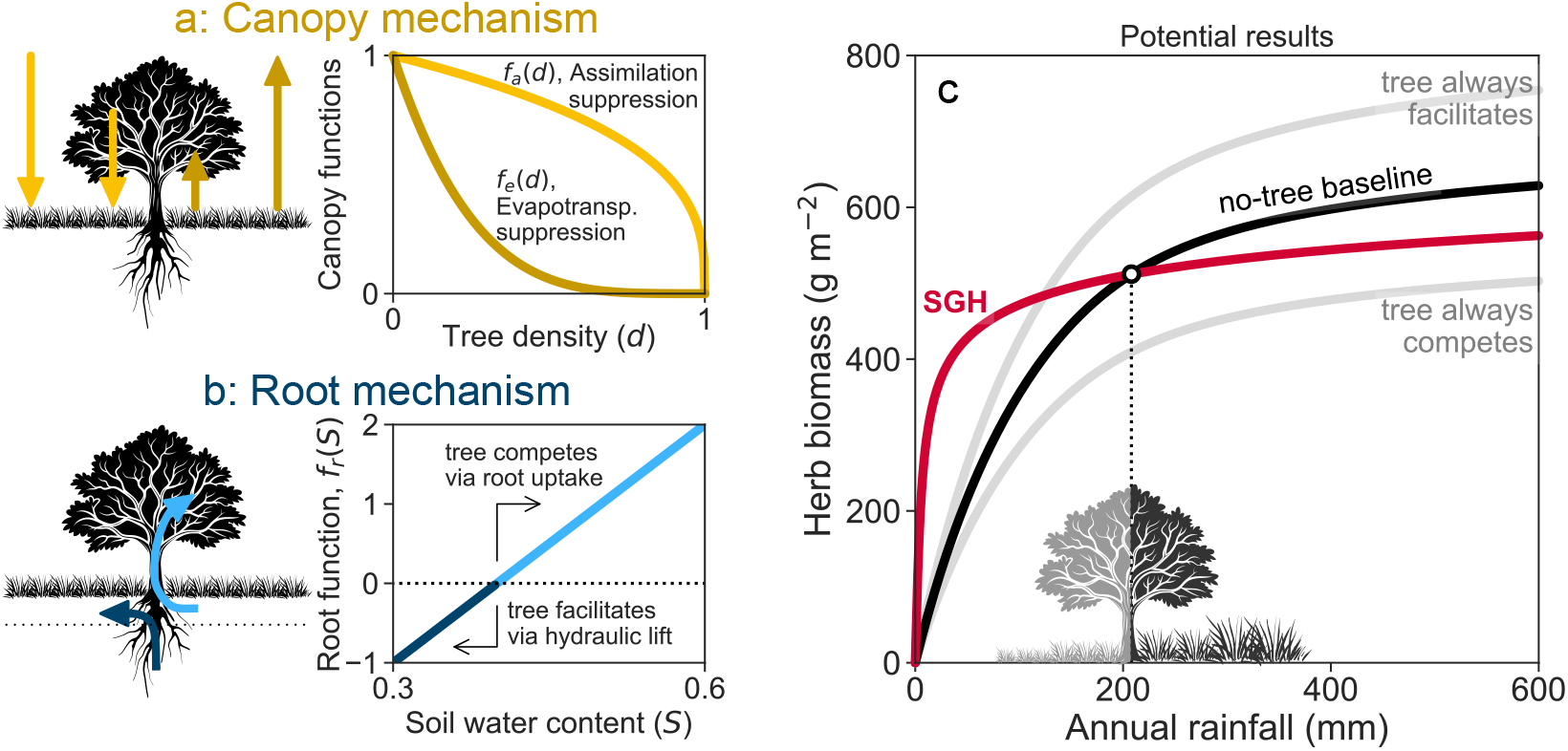
Assumptions of the two submodels (a, b) and potential model outcomes (c). (a) The canopy mechanism assumes that tree density suppresses both photosynthetic assimilation (yellow) and water loss by evapotranspiration (brown). However, the reduction in evapotranspiration with increasing tree density is greater than the reduction in photosynthesis, which is a necessary condition for facilitation under shading. (b) The root mechanism focuses on water uptake (light blue) and hydraulic lift (dark blue), where water from deep layers is transported upward by the tree and increases moisture in shallow soil. The curve reflects the net effect of these two opposing processes: when the soil is dry, steep water potential gradients between deep and shallow layers favor upward water movement throught the roots, producing negative net uptake (a gain to the upper soil). As soil moisture increases, water extraction by roots becomes more efficient, and uptake outweighs hydraulic lift, resulting in higher (more positive) net uptake values. (c) Potential model outcomes are illustrated by comparing rainfall–biomass relationships with and without trees (black line). In the two simple scenarios (grey), trees either consistently facilitate or consistently inhibit herb growth across the rainfall gradient, whereas the Stress Gradient Hypothesis (SGH; red line) predicts a transition from facilitation to competition, marked by the intersection between the black and the red lines.

## 2 Results

We developed a consumer-resource model that describes the coupled dynamics of herbs’ biomass and soil moisture as follows:

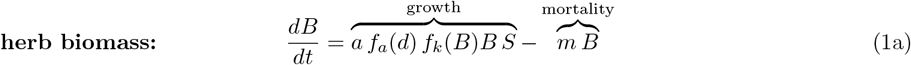

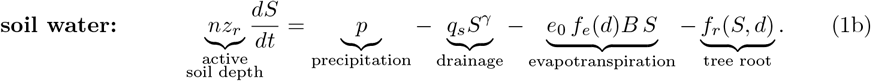

The first equation tracks the change in herb biomass (*B*) over time, which is governed by growth and mortality processes. The second equation represents the dynamics of soil moisture (*S*), which is influenced by gains from precipitation (*p*) and losses due to drainage, evapotranspiration, and a tree-root effect. Due to their much slower dynamics, trees are represented as a constant parameter for tree density (*d* representing canopy cover or root density). This parameter affects both the canopy-suppression factors on herb growth (*f*_*a*_(*d*)) and on evapotranspiration (*f*_*e*_(*d*)), as depicted in Fig. 1a. The impact of the root mechanism *f*_*r*_(*S, d*) on soil water is depicted in Fig. 1b. We further assumed that when water and light are ample, other factors such as nutrients or genetic limits constrain herb growth, represented by a carrying capacity term *f*_*k*_(*B*) = 1 − *B/k*. For simplicity, we investigated each mechanism separately: when examining canopy effects, we removed root effects, and vice versa.

When water is the main limiting factor and tree density is constant, our model, like previous mechanistic models ^36;37^, shows that the interaction between trees and herbs remains either facilitative or competitive along the entire precipitation gradient. However, a key finding is that introducing a new limiting factor (*f*_*k*_(*B*) in the model), which can represent nutrient limitation, or a genetic size limit, is a **necessary condition** for the Stress Gradient Hypothesis transition to occur (see Supplementary Section S1). When this condition is met, both the canopy and root mechanisms can produce a clear shift from facilitation to competition as precipitation increases. The following explores how each of these mechanisms drives this transition.

In the **canopy mechanism**, in the absence of trees, herb biomass increases with precipitation, showing a saturation pattern as the curve’s slope decreases (black line in Fig. 2a). With some trees (light curve), herb biomass is higher than the no-tree baseline at low precipitation but is reduced at higher precipitation levels, showing a clear transition from facilitation to competition. At very high tree density (dark curve), herb biomass is suppressed across the entire precipitation gradient.

**Figure 2.**
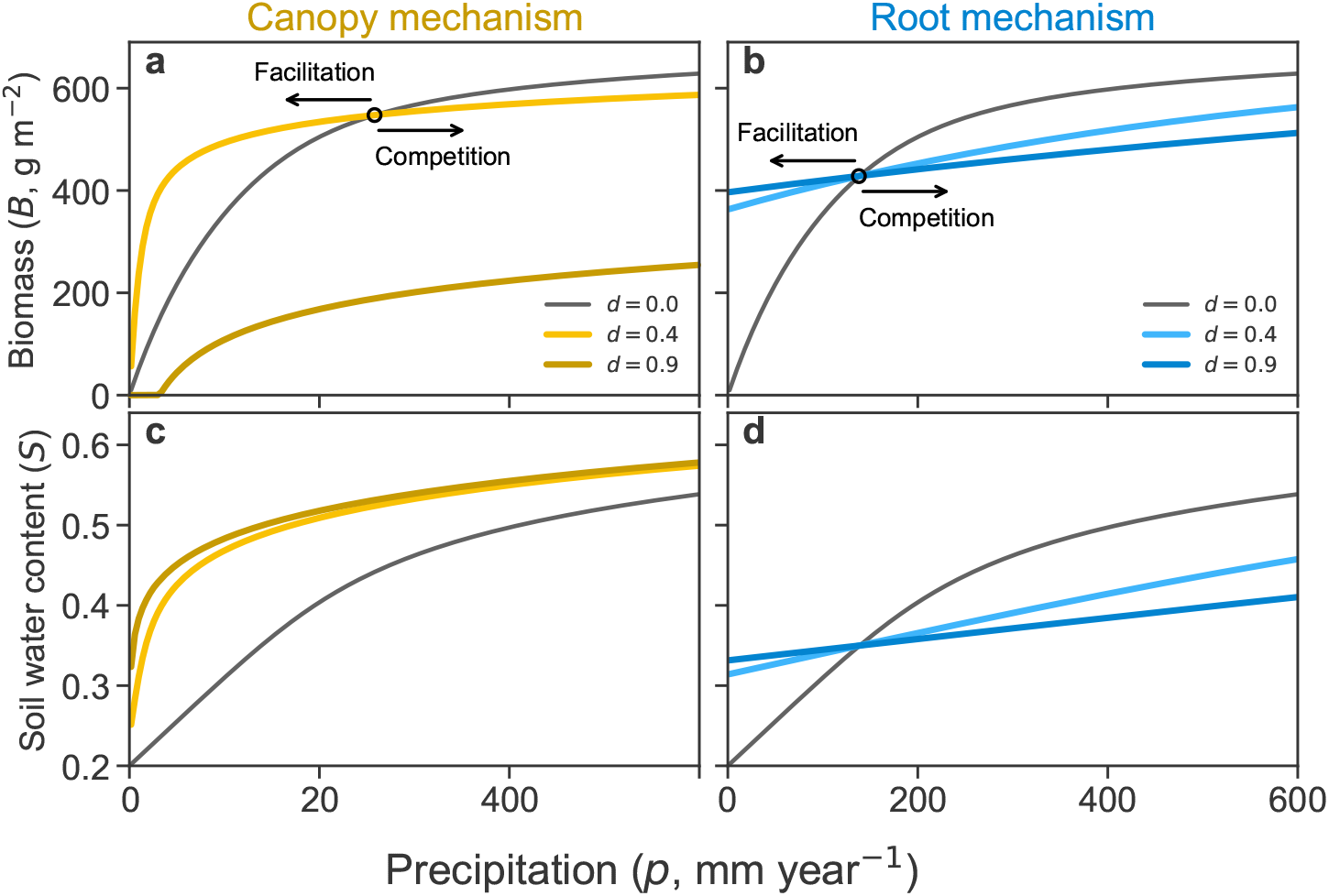
The transition from facilitation to competition is produced by two different mechanisms. Herb biomass and soil water solutions of (Eqs. (1)) are shown for the canopy mechanism (panels a,c) and the root mechanism (panels b,d). Black lines give the no-tree baseline (*d* = 0), light solid curves show moderate tree density (*d* = 0.4) and dark solid curves high tree density (*d* = 0.9). Facilitation (competition) occurs when the herb biomass solid curves are above (below) the baseline curve. Other parameter values are as reported in Table 1.

This pattern is a result of a tug of war between two opposing forces exerted by the trees. First, trees facilitate growth by providing shade, which reduces evapotranspiration and conserves soil water. This effect is represented by the function *f*_*e*_(*d*), leading to greater soil water availability relative to a no-tree environment (see Fig. 2c). Second, trees inhibit herb’s growth by reducing light availability through the function *f*_*a*_(*d*). This factor, along with other limiting elements like nutrients or grazing (captured by the carrying capacity term, *f*_*k*_(*B*)), down-regulates herb assimilation. The combined effect is represented by the product *f*_*a*_(*d*)*f*_*k*_(*B*).

**Table 1.** Variables and parameters with typical value ranges.

| Variables |  |  |  |
| --- | --- | --- | --- |
| Symbol | Units | Values | Description |
| $B$ | $\text{g m}^{-2}$ | 0–1000 | Herb biomass density |
| $S$ | — | 0–1 | Relative soil water content |
| Parameters |  |  |  |
| $a$ | $\text{d}^{-1}$ | 0.05 | Assimilation rate |
| $\beta_a$ | — | 1/3 | Shading function exponent |
| $\beta_e$ | — | 4 | Shading function exponent |
| $k$ | $\text{g m}^{-2}$ | 1000 | Carrying capacity |
| $m$ | $\text{d}^{-1}$ | 0.01 | Mortality rate |
| $p$ | $\text{mm y}^{-1}$ | 0–600 | Precipitation rate |
| $e_0$ | $\text{mm d}^{-1}/\text{g m}^{-2}$ | 0.005 | Max evapotranspiration rate per unit biomass density |
| $q_{sat}$ | $\text{mm d}^{-1}$ | 800 | Saturated hydraulic conductivity |
| $\gamma$ | — | 10 | Deep infiltration exponent |
| $n$ | — | 0.4 | Soil porosity |
| $z_r$ | mm | 300 | Herb rooting depth |
| $d$ | — | 0–1 | Woody plant density |
| $S_h$ | — | 0.3 | Hygroscopic point |
| $S_{fc}$ | — | 0.6 | Field capacity |
| $\lambda_h$ | $\text{mm d}^{-1}$ | –2 | Max tree water provision |
| $\lambda_{fc}$ | $\text{mm d}^{-1}$ | 10 | Max tree water usage |

**Note.** Variable $B$ range derived from Mussery et al.<sup>64</sup>. Variable $S$ range and parameters $q_{sat}$ , $\gamma$ , $n$ , $z_r$ , $S_h$ , $S_{fc}$ derived from Rodriguez-Iturbe et al.<sup>60</sup>. Parameter $a$ derived from James and Drenovsky<sup>65</sup>. The ratio between $\beta_a$ and $\beta_e$ is derived from Pons et al.<sup>66</sup>. Parameter $p$ range derived from Huang et al.<sup>67</sup>. Parameter $\lambda_{fc}$ is derived from Shiferaw et al.<sup>68</sup>. Parameter $e_0$ is derived from Allen et al.<sup>69</sup>; Garnier et al.<sup>70</sup>.

At low precipitation, herb biomass is low, so the carrying capacity term *f*_*k*_(*B*) is very weak (close to 1). In this dry scenario, the tug of war is mainly between the water-saving benefit of *f*_*e*_(*d*) and the light reduction effect of *f*_*a*_(*d*). When tree density is low, the extra soil water overpowers the minor loss in light, facilitating herb growth (light curve in Fig. 2a). However, when tree cover is high, the reduction in light becomes too strong, suppressing the herbs (dark curve). As precipitation increases, herb biomass also rises. This strengthens the carrying capacity term *f*_*k*_(*B*) (making it closer to zero) and tips the balance. The combined competitive effect of reduced light and carrying capacity (*f*_*a*_(*d*)*f*_*k*_(*B*)) becomes stronger than the facilitative effect of water conservation. In other words, under high rainfall, water is no longer the limiting factor, and therefore, the benefit of reduced water loss is not enough to compensate for the reduction in photosynthesis, leading to a transition from net facilitation to net competition.

For the **root mechanism**, all non-zero tree densities show a similar transition pattern: herb biomass is higher than the baseline at low precipitation and lower at high precipitation (see Fig. 2b). As tree density increases, both effects intensify along the precipitation gradient. This is because the tug of war between facilitation and competition is expressed by a single function, *f*_*r*_(*S, d*), which represents the net effect of tree roots on soil water available in the herb root zone.

This function captures the tipping of the balance as soil water content changes. At low soil water levels, tree roots can lift water from deeper soil layers to the herb root zone, facilitating growth. As soil water content increases, hydraulic lift is no longer possible. Beyond this point, roots begin to compete with herbs by taking up water from the same soil layer. Counterintuitively, the root term *f*_*r*_(*S, d*) by itself does not produce a facilitation-to-competition switch (see Supplementary Section S1). With water as the only limiting resource (i.e., *f*_*k*_(*B*) = 1), the equilibrium soil moisture *S*^*⋆*^ is set by the consumer and is independent of the precipitation supply *p* (R* logic; ^33^). Because *f*_*r*_ acts through *S* rather than directly through *p*, the sign of the interaction is fixed by whether *S*^*⋆*^ lies below or above the hydraulic switching range, so it does not change along the rainfall gradient. However, when a second growth constraint is introduced through the carrying-capacity term *f*_*k*_(*B*), the outcome changes: as *p* increases, biomass approaches its limit and cannot deplete water further, so *S*^*⋆*^ rises. This upward shift in *S*^*⋆*^ carries the system across the hydraulic threshold, turning facilitative lift into competitive uptake and yielding the observed transition. Put simply, *f*_*k*_(*B*) caps biomass at high rainfall, weakening consumption and allowing soil water to accumulate, which moves *S*^*⋆*^ into the competitive domain of *f*_*r*_(*S, d*).

Notably, the two mechanisms produce very different biomass responses at low precipitation. In the canopy mechanism, precipitation is the only water source, so all curves must start from the origin, zero biomass at zero rainfall. A small initial increase in precipitation leads to stronger facilitation, which is visible as a widening gap between the low-tree-density curve and the no-tree baseline (light and black curves in Fig. 2a, respectively). This facilitative gap eventually narrows before disappearing at the transition to competition. In contrast, the root mechanism includes an additional water source: hydraulic lift from deeper soil layers during dry surface conditions. This allows herb biomass to persist even without precipitation. The gap between tree-density curves and the baseline shrinks steadily as precipitation increases, until it reaches the transition point. This distinct pattern in the biomass gap is key to understanding the contrasting responses in interaction intensity between the two mechanisms.

For the canopy mechanism, the interaction intensity based on the absolute difference is unimodal (see Fig. 3a). This pattern directly results from the widening and eventual vanishing of the gap between the biomass curve with trees and the no-tree baseline curve, as previously discussed. However, when using the relative log response ratio (see Fig. 3c), the interaction intensity decreases monotonically for *p* below the transition threshold. In contrast, the root mechanism yields monotonically-decreasing positive interaction intensities, regardless of whether absolute difference or relative log response ratio is used. A broader discussion of the canopy mechanism’s interaction intensity in the full (*p, d*) parameter space is given in Supplementary Section S2.

**Figure 3.**
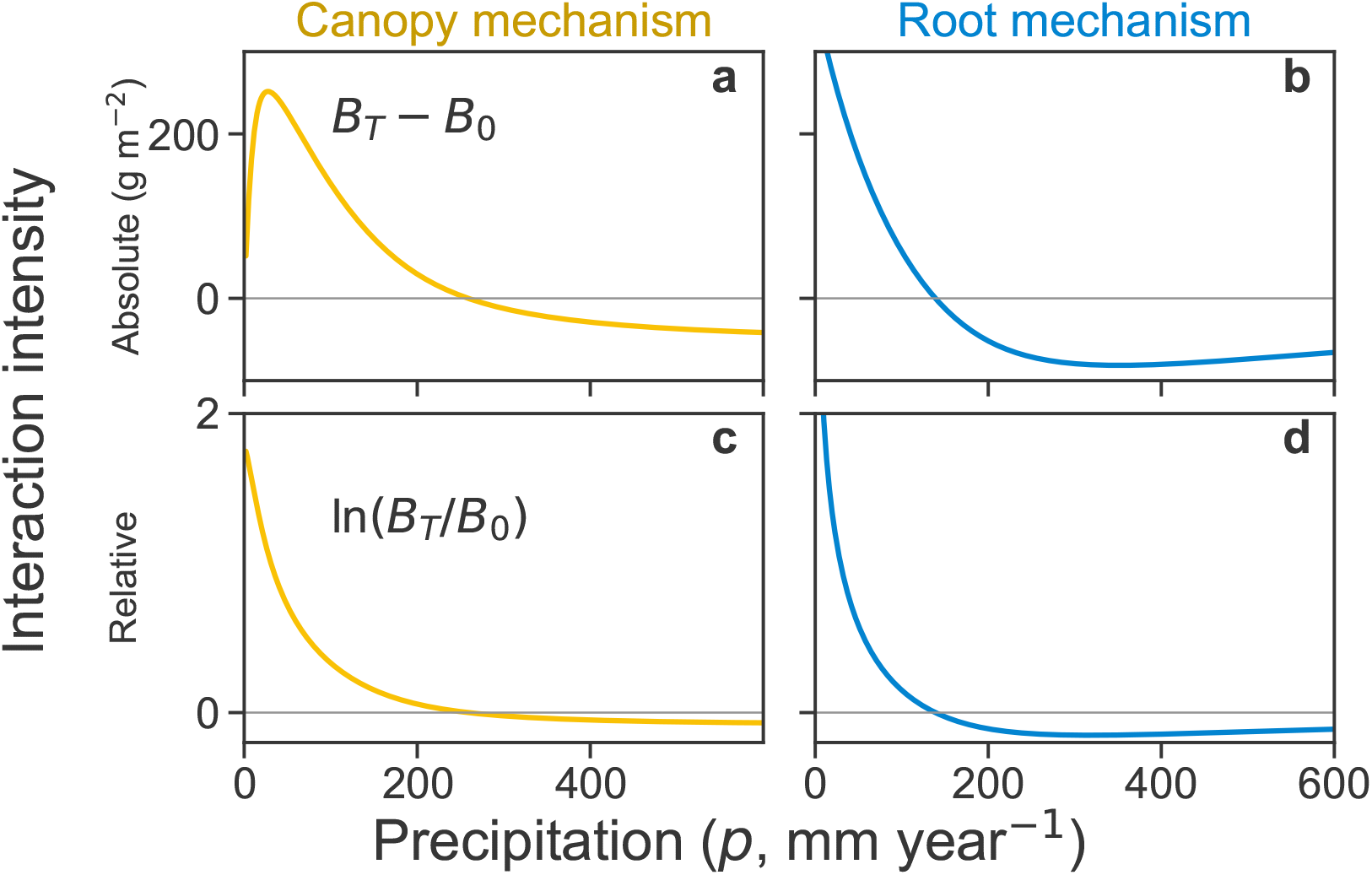
Interaction intensity patterns depend on both mechanism and metric. The gray lines represent zero interaction. Positive interactions indicate facilitation by trees; negative interactions indicate competition. Panels a and b show the absolute interaction intensity, *B*_*T*_ *− B*_0_, where *B*_0_ and *B*_*T*_ denote herb biomass in the absence and presence of tree density, respectively. Panels c and d show the relative log response ratio, ln(*B*_*T*_ */B*_0_). Parameters: *d* = 0.0 and *d* = 0.3 were used to compute *B*_0_ and *B*_*T*_, respectively; other parameters are given in Table 1.

## 3 Discussion

The Stress Gradient Hypothesis has guided decades of work but has largely been articulated in verbal or phenomenological terms ^9;19;20;26;27;29;30^. We show that a minimal consumer-resource model linking canopy shading or root water redistribution to herb biomass generates the observed shift from facilitation to competition along rainfall gradients. Our model also specified when the shift should and should not arise. Finally, the framework also clarifies the expectation on interaction intensity, with a mid-gradient peak in the facilitation under shading and a monotonic decline from positive to negative effect under the root pathway.

Our aim is to provide general insight and testable predictions rather than site-specific accurate description. Accordingly, the model is intentionally simple yet mechanistic: water balance is explicit, and interaction signs emerge from the equations rather than being imposed a priori. This simplicity enables a thorough understanding of each parameter (See Supplementary Section S3).

The model focuses on the primary pathways through which woody plants influence herbs: canopy shading and root water redistribution. It keeps tree density fixed while examining steady-state contrasts. Other processes, such as grazing protection, nutrient supply, or spatial patterns ^31;36–38^ can be incorporated where they are expected to matter (see the Supplementary Sections S4 and S8 for transient dynamics and varying tree density). We therefore view this framework as a baseline for systems where water and light competition dominate, and as a foundation for targeted extensions that incorporate additional mechanisms.

The stress gradient hypotheses have been invoked for many stress types ^9;18–23^. Yet, as Maestre et al. ^29^ noted, “Stress is not a precise concept, and therefore it is difficult to apply quantitatively to communities or ecosystems”. Here we focus on mechanisms tied directly to plant water balance, while recognizing that other stressors such as freezing, toxicity, or salinity, are likely to require different model structures. The same framework can also be applied to evaporative demand, another driver of aridity, which is influenced by temperature and humidity. This driver can change the magnitude of facilitation and competition and shift the transition point along the gradient. Still, the model consistently predicts a shift from net facilitation to net competition as water limitation relaxes (see Figure in Supplementary Section S5).

The model resolves the empirical question whether stress should be represented by resource supply rate (precipitation) or by resource abundance (soil water content) ^9;36;37;39^. It shows that precipitation or evaporative demand are the appropriate measures for quantifying water stress in such systems (see Supplementary Section S6). Soil water content, in contrast, is not an independent driver but an emergent outcome of interacting processes including precipitation, evapotranspiration, biotic uptake, and substrate properties ^9;36;39^. Treating soil water as externally fixed cuts off these mechanistic links and removes the causal connection between resource supply and species interactions. Only when stress is parameterized as a supply rate can the facilitation–competition interplay characteristic of the SGH be captured ^9;36;37;39^.

### 3.1 Canopy mechanism

Shading is a ubiquitous factor plant growth and community dynamics ^40–43^. This mechanism is central in conceptual models that generate the transition from facilitation to competition ^9;20^. Our findings indicate that shading can simultaneously enhance and constrain growth, with implications far beyond tree–herb interactions.

In the model, shading acts solely by reducing light, so any reduction in radiation, whether caused from trees, slope aspect, buildings, or solar panels, should generate similar responses. In dry conditions, shaded hillslopes, north-facing in the northern hemisphere and the reverse in the southern hemisphere, should be more productive than sun-exposed slopes. In wet conditions, the pattern should reverse. Variation with slope aspect is a long-standing observation in botany ^44;45^, yet we are not aware of a mechanistic model that explains this global pattern. The same logic applies to urban ecosystems, where buildings cast shade, and to agricultural settings, with the co-location of solar panels and crops (agrivoltaics) being on the rise over the last two decades ^46^.In such settings, the model predicts when agrivoltaics will increase or decrease productivity depending on environmental conditions

The model highlights several conditions that have seldom been investigated in empirical studies along gradients. First, a necessary condition for facilitation under shading is that the proportional reduction in evapotranspiration with increasing shade exceeds the proportional reduction in photosynthesis. Although this likely holds for many plants, there are clear exceptions, such as species that require high light and are sensitive to shade. We therefore suggest that empirical tests of the shading mechanism begin by verifying this assumption.

For the transition from facilitation to competition to occur without a change in tree cover, another condition must be met. Under high rainfall, water must cease to be the main limiting factor (the carrying capacity effect). This implies that a qualitative switch is less likely when moving from an arid to a semiarid system if both remain water-limited. Instead, a switch is expected only when crossing into a system limited by another resource, such as nutrients (see Supplementary Section S7, which shows that carrying capacity is equivalent to an additional essential resource). This result may explain empirical studies that do not observe a shift along precipitation gradients ^17;29;47;48^ and underscores that the transition should not be viewed as inevitable.

Alternatively, shading can lead to a facilitation-to-competition transition without introducing carrying capacity, but only when tree density increases with rainfall (See Supplementary Section S8). This occurs because at high tree cover, light becomes the primary limiting factor and offsets the positive effects of shading on water balance. This density effect can be further enhanced by photoinhibition, where excessive light inhibits photosynthesis. We therefore recommend that future empirical studies quantify how tree density changes along the gradient and manipulate tree cover directly, or mimic shading with shade cloth. Such an approach is necessary to determine whether interaction outcomes change under a constant shade level (Fig. 2), or arise from shifts in canopy density along the gradient (Supplementary Section S8).

### 3.2 Root mechanisms

Root-mediated effects in natural settings can be highly variable because they depend on root architecture and soil properties ^11;49^. In the canopy mechanism, we used a two-layer simplification in which deep tree roots do not change the soil water available to herbs ^50^. By contrast, in the root mechanism, we assumed partial overlap in rooting depth so that trees and herbs draw from the same near-surface water. We further assumed hydraulic lift, whereby trees move water upward from deeper layers when the surface is dry and draw water from the herb layer when it is wet.

When trees increase soil moisture under dry conditions and reduce it under wet conditions, a transition from facilitation to competition can occur. Although we initially expected this transition to arise inevitably from the root mechanism, we found that it occurs only when another factor limits biomass at high rainfall; otherwise, equilibrium soil moisture remains constant along the gradient. Notably, although there is empirical evidence for hydraulic lift ^13^, its strength and prevalence remain uncertain, so we treat it as secondary to shading for broad-scale patterns. Alternatively, a more common way trees may increase soil moisture is by enhancing infiltration ^11^. While infiltration alone does not cause a facilitation-to-competition transition ^37^, our model shows that it can do so when combined with shading: under low precipitation, the positive effects of increased infiltration dominate, whereas at high rainfall, the negative effects of shading prevail (Supplementary Section S9).

### 3.3 Concluding Remarks

Our findings help reconcile conflicting reports on interaction strength along rainfall gradients by showing that the expected pattern depends first on the mechanism. Under the canopy pathway, the absolute difference in biomass is hump-shaped, with maximum facilitation at intermediate rainfall, whereas under the root mechanism, the interaction declines monotonically from positive to negative. A second source of variation is the metric, and this sensitivity applies to shading in particular: only the absolute measure yields a unimodal pattern, whereas relative measures decline with rainfall, consistent with earlier suggestions ^9^.

Importantly, tree density further modulates these patterns of interactions along aridity gradients (Supplementary Section S2). Hence, empirical patterns can only be interpreted accurately when tree density is quantified. Under low precipitation, facilitation peaks at intermediate density, while low and high tree densities weaken it. As precipitation increases, progressively lower densities are sufficient to shift the balance from facilitation to competition. Eventually, at high rainfall, even near-zero tree density reduces herb performance, so further changes in density no longer cause a qualitative shift. ^9^.

Many empirical studies report a shift from facilitation to competition ^14–16^, and many do not ^9;28–31^. In the lens of our framework, cases without a shift arise when (i) other pathways dominate, for example, protection from herbivory, or (ii) when the conditions for a shift are not met, for example, when shade suppresses photosynthesis more than evapotranspiration. This perspective moves the discussion from whether the hypothesis holds to which mechanism operates. It also points to practical tests, pairing shade manipulations with canopy and soil water measurements, and reporting both absolute and proportional changes.

Looking ahead, climate change is exacerbating water limitations in many regions ^51–54^, while woody cover is changing due to drought, fire, shrub encroachment, afforestation, and altered land use ^1;2;5–7^. A compact mechanistic framework can help anticipate where shade will enhance herb production by conserving water and where it will suppress production due to light limitations, and it can guide restoration and conservation efforts toward interventions that match local mechanisms. By building on the Stress Gradient Hypothesis and giving it simple, testable conditions, this framework connects a classic idea to actionable predictions for conserving dryland ecosystems in a rapidly changing world.

## 4 Methods

We developed a consumer-resource model (see Eq. (1)) that describes the dynamics of herbaceous biomass density (*B*, kg m^*−*2^) ^36;37^, and relative soil-water content (*S*, dimensionless) ^55^.

The model makes three key assumptions: (i) Tree biomass changes on a much longer time scale than herb biomass, so tree density is treated as constant and remains in quasi steady state relative to the herbs and soil water. (ii) Herb roots occupy only the upper soil layer, whereas tree roots also reach deeper layers, allowing trees to lift water upward or to draw water away from the herb rooting zone; (iii) Herb growth is limited by light and by soil moisture, yet only water can accumulate over time and therefore is described by a balance equation. To incorporate constraints on growth beyond water and light, we include a carrying capacity term, which is equivalent to assuming another limiting resource, such as an essential nutrient (see Supplementary Section S7).

The dynamics of herbaceous biomass density *B* are governed by growth and mortality terms (Eq. (1a)). The dynamics of relative soil-water content *S* are dictated by a water-balance equation ^55^, whose input is precipitation and whose outputs are drainage to deeper soil layers and evapotranspiration, while tree-root processes can function as both inputs and outputs depending on direction of water flow.(Eq. (1b)). Herb biomass and water are averaged over the horizontal dimensions, and water is averaged over the active soil depth *nz*_*r*_, following a traditional bucket-model approach ^56^.

The functions *f*_*a*_(*d*), *f*_*k*_(*B*), *f*_*e*_(*d*), *f*_*r*_(*S, d*) are modular components of the model that can be turned on or off, either when switching between model variants or when testing the impact of different limiting factors. The features that are common to all realizations of the model are: (i): Herb biomass growth rate is linearly dependent on soil water content *S* (when the carrying capcity term is negligible). (ii) Herb biomass mortality rate is proportional to biomass. (iii) Evapotranspiration is down-regulated by soil moisture availability, following a linear *β* function of *S* ^57;58^, and is proportional to herb biomass density. (iv) Drainage is modeled by a highly nonlinear function of soil moisture ^59^, commonly used in ecohydrological modeling ^60^. (v) The logistic growth term *f*_*k*_(*B*) = 1 *− B/k* was used throughout this paper and was only turned off (*f*_*k*_(*B*) = 1) in Supplementary Sections S1 and S8, where we studied the effects of removing additional limiting growth-limiting factors beyond water and light. (vi) All the model parameters (Table 1) are constant. In particular, precipitation rate *p* is understood as the total precipitation of the growing season divided by its duration; it is reported in mm y^*−*1^ instead of mm d^*−*1^ throughout the paper to enhance interpretability.

Our model belongs to the family of coupled soil moisture and biomass models developed for other purposes ^61–63^. The key difference is that many earlier models treated biomass growth and evapotranspiration as the same function scaled by a constant conversion factor, water use efficiency, defined as water loss per unit carbon assimilation. Here, both processes depend on moisture, biomass, and tree density, but they have different functional forms, so water use efficiency varies across environments. This follows from the fact that shading can affect evapotranspiration and assimilation differently (Fig. 1a). In addition, relative biomass growth declines with size through the logistic term, whereas evapotranspiration remains proportional to biomass. These assumptions are both biologically plausible and necessary for the mechanisms we study: if water use efficiency were constant across shading levels, the trivial pattern where partial shade is beneficial but heavy shade is detrimental could not arise.

Below, we discuss in more detail the two main mechanisms employed in this paper.

### Canopy mechanism

Shading affects both plant growth, by reducing light availability and thus photosynthesis, and transpiration, by lowering temperature and radiation levels, which helps retain soil moisture and improve water availability for herbs. When the shading mechanism is “on”, the root mechanism is disabled by setting *f*_*h*_(*S, d*) = 0, implying a complete partitioning of the soil into two distinct niches, the top available for herbs only, and deeper layers accessible to trees only.

Biomass growth is down-regulated by shading via *f*_*a*_(*d*), while evapotranspiration is down-regulated via *f*_*e*_(*d*). The shading functions are 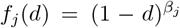, where *j* = *{a, e}*. Figure 1a shows the nonlinear decline of these functions with tree density. A necessary condition for facilitation is *β*_*e*_ *> β*_*a*_; assimilation is therefore less inhibited by shade than transpiration.

### Root mechanism

Trees can influence herbs access to water in two ways: they can lift water from deeper layers into the herbs rooting zone when surface soil is dry (facilitation), and they can uptake water from that zone (competition). The combined effects of hydraulic lift and tree water uptake is described by the function *f*_*r*_(*S, d*). When the root mechanism is “on”, the shading mechanism is disabled by setting *f*_*a*_(*d*) = *f*_*e*_(*d*) = 1. The expression for the root function reads:

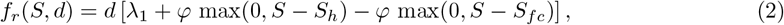

where the slope *φ* = (*λ*_*fc*_ *− λ*_*h*_)*/*(*S*_*fc*_ *− S*_*h*_). For *S* smaller than the hygroscopic point *S*_*h*_, the upper soil is too dry for tree roots to uptake water, and hydraulic lift reaches its maximum ability to bring water from deeper soil layers to the topsoil (*λ*_*h*_). For *S* greater than the field capacity *S*_*fc*_, trees cease to benefit from increasing soil water content, and uptake water at a rate *λ*_*fc*_. Between *S*_*h*_ and *S*_*fc*_, the root function varies linearly between *λ*_*h*_ and *λ*_*fc*_. Regardless of soil water content, the root function depends linearly on tree density *d*. Figure 1b shows the function *f*_*r*_ in the range *S*_*h*_ *< S < S*_*fc*_, for zero tree density (*d* = 0) and maximal tree density (*d* = 1).

### 4.1 Numerical solutions

Numerical analyses were performed with Python 3.12, using the libraries NumPy 2.0 and SciPy 1.13. Steady-state solutions (*dB/dt* = *dS/dt* = 0) were obtained by finding the roots of the right-hand side of Eqs. (1) with scipy.optimize.fsolve, using random starting estimates for the roots (0.5 *< B <* 1.5 kg m^*−*2^ and 0.5 *< S <* 0.6). A root is accepted when *B >* 0.01 kg m^*−*2^ and 0 *< S <* 1; if these criteria are not met, a new set of random starting estimates is chosen. This procedure is repeated up to 10 times, after which the steady-state solutions are estimated by numerically integrating Eqs. (1) with scipy.integrate.solve_ivp (stiff solver, default tolerances) up to a final time of 10 thousand days, and the final configuration is taken as the steady-state solutions.

## 4.2 Code Availability

The code to run the model can be found in the following Zenodo repository https://doi.org/10.5281/zenodo.17641468.

## 4.3 Model Parameters

The model variables and parameters are summarized in Table 1. The specific parameter values (or their ranges) were chosen from typical values found in the literature.

## 5 Acknowledgments

We thank Barel Tsafon, Atay Mor, Santanu Das, Erez Feuer and Moshe Shachack for comments on earlier drafts. We thank Jonathan Friedman for useful discussion. Figure 1 utilized the images: [oak by Levi](https://thenounproject.com/browse/icons/term/oak/), roots by wahab marhaban, [grass by Jhonatan](https://thenounproject.com/browse/icons/term/grass/), all from the Noun Project (CC BY 3.0). This work was supported by the Center for Sustainability, and by the Research Center for Agriculture, Environment and Natural Resources, both of the Hebrew University of Jerusalem.

## 7 Author Contribution

Conceptualization: O.H., Y.M., and N.D.; Analysis: O.H. and Y.M.; Funding acquisition: Y.M. and N.D.; Supervision: Y.M. and N.D.; Visualization: O.H. and Y.M.; Writing—original draft: O.H.; Writing—review and editing: Y.M. and N.D.

## Supplementary Information

### S1 Role of carrying capacity in the SGH patterns

We explore here the model results for each of the two mechanisms in the case where the logistic growth term is rendered inactive. This is achieved by taking the carrying capacity *k → ∞*, which makes *f*_*k*_(*B*) = (1 − *B/k*) → 1. We can now find analytical expressions for the steady-state solutions of the model described in Equations (1) in the main paper. The nontrivial (*B* ≠ 0) steady-state solutions (*B*^*⋆*^, *S*^*⋆*^) read:

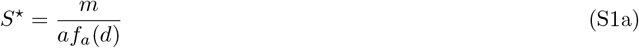

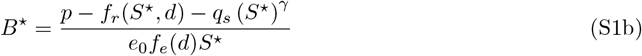

These solutions are a typical instance of the R-star rule ^1^ and from them we can express two important features:

1. The steady-state resource level *S*^*⋆*^ (generally called *R*^*⋆*^ in consumer-resource modeling, therefore the name of the rule) is independent of the resource supply rate, *p*.
2. The nontrivial (*B≠* 0) steady-state consumer level *B*^*⋆*^ is linear in the resource input rate, *p*.

Figure S1 shows the steady-state biomass *B*^*⋆*^ (top panels) and relative soil water content *S*^*⋆*^ (bottom panels). The left and right columns correspond to the shading mechanism and the root mechanism, respectively. A careful examination of the steady-state solutions (S1) teaches us that:

**For the shading mechanism**, 0 ≤ *f*_*e*_(*d*) *< f*_*a*_(*d*) *<* 1 for any *d >* 0 (see inset in Fig. S1c). From these facts it follows that the linear function for the positive tree-density solution (*B*^*⋆*^(*p, d >* 0)) has a higher slope and a lower intercept^2^ than the no-tree solution (*B*^*⋆*^(*p, d* = 0)). Because of its lower intercept, the positive density solution surpasses the zero-tree density solution from below (lower *B*). This means that eliminating the logistic growth term from the model **reverses the order of the transition** described by the SGH: we have competition for *p* below the transition point, and facilitation for *p* above that point. Furthermore, for the parameter values we used here, the transition point occurs at extremely low values of *p* and *B*. Looking from ‘farther away’, the interactions appear facilitative for most of the precipitation gradient (Fig. S1a).

**Figure S1.**
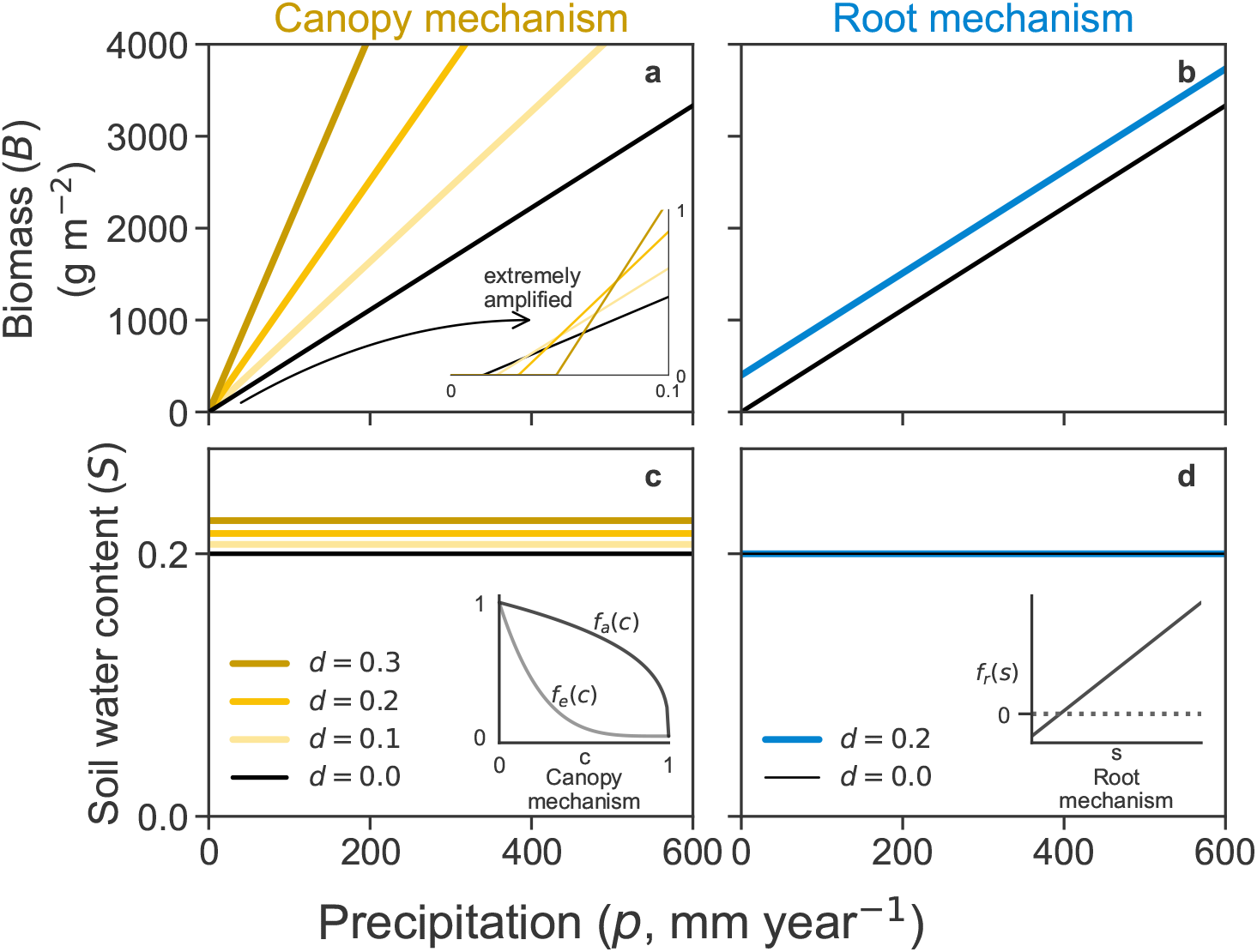
Steady-state biomass (panels a,b) and soil water content (panels c,d) solutions as functions of precipitation. The left column corresponds to the shading mechanism, and the right column corresponds to the root mechanism. The inset in panel a shows an extreme amplification in *p* and *B* of the solutions, emphasizing the intersection points between the no-tree solution (*d* = 0) and the other solutions with *d >* 0. The insets in panels c and d are provided as reminders of the relevant functions of each mechanism.

**For the root mechanism**, two straight-line solutions *B*^*⋆*^ always have the same slope, regardless of their *d* value, and therefore never intersect. The root mechanism can be either facilitative or competitive, depending on the soil water content (see inset in Fig. S1d). Figure S1b shows a scenario where the interaction is always facilitative. By changing the model parameters *m, a, d* (see Eq. S1a), one can also find higher soil water content levels that always yield competitive interactions.

The arguments above demonstrate that, whenever tree density is constant, the **introduction of an additional growth-limiting factor** that strengthens with increasing precipitationis a **necessary condition** for the emergence of the transition described by the SGH.

### S2 Interaction intensity

As a rule, trees facilitate herb growth when their density is low and the system is under water stress (low *p*). Competition over water arises at high tree density values and under low water stress. Focusing on the canopy mechanism, Figure S2 shows that in the (*p, d*) parameter space, the boundary between facilitation and competition (solid black curve) follows a negative relation: as precipitation increases, a lower tree density is sufficient to shift the balance from facilitation to competition. This border is the same for both interaction intensities definitions, since zero interaction intensity means that the biomass solution with tree density (*B*_*T*_) equals the biomass solution with no tree density (*B*_0_). The area hatched in black at the top of both panels, where tree density is very high, indicates the region in the parameter space where there are only trivial (*B* = 0) vegetated solutions.

**Figure S2.**
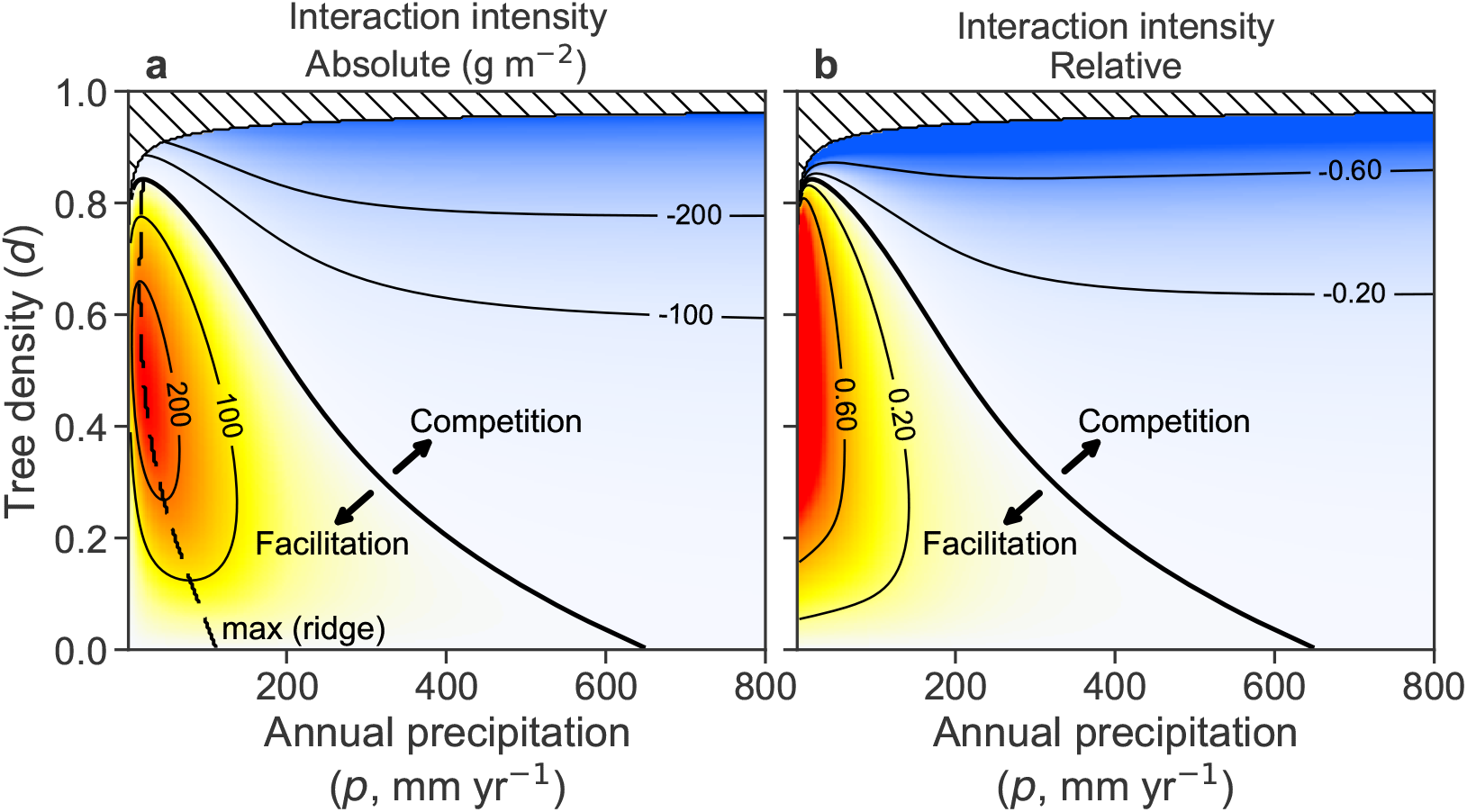
Interaction intensity varies jointly with precipitation and tree density. Interaction intensity in the canopy submodel is shown as a function of annual precipitation (*p*) and tree density (*d*), measured as (left) the absolute difference in herb biomass, *B*_*T*_ *− B*_0_, and (right) the relative log response ratio, ln(*B*_*T*_ */B*_0_), where *B*_0_ and *B*_*T*_ denote herb biomass in the absence and presence of trees, respectively. Purple shades indicate facilitation (positive values) and orange shades indicate competition (negative values). Both metrics reveal a transition (black line) from facilitation at low rainfall and tree density to competition at high rainfall and tree density. The dashed black line on the left panel indicates the locus of maximum facilitation for a fixed tree density value [ 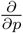 (int. intensity) = 0], for the absolute interaction intensity.

As shown in Fig. 3a, we can get unimodal curves when using the absolute interaction intensity. The dashed black curve in Fig. S2a shows the location of the local maxima in the (*p, d*) parameter space. To the left of this curve, the interaction intensity is positive and increases with precipitation. Once we cross this curve (from left to right), interaction intensity is still positive, but it decreases with precipitation. This ridge-like curve disappears when we use the relative interaction intensity instead: there is only monotonic decrease with precipitation.

### S3 A deeper look into the model

In order to find useful ways of thinking about the model, we perform a non-dimensionalization of the equations. For our purposes, it makes more sense to describe the second equation as the rate of change of absolute soil water content *W* = *nz*_*r*_*S*, instead of the equivalent *nz*_*r*_ *dS/dt* as shown in Eq. (1). Since the active soil layer *nz*_*r*_ has length dimension (*z*_*r*_ is the depth of the herb rooting zone, while the porosity *n* is non-dimensional), absolute soil water content *W* is also a length (it is commonly called soil water depth). Finally, we note that 1 mm of water is equivalent to 1 L m^*−*2^, so reporting *W* as length is the same as volume per unit area. The full model equations now read:

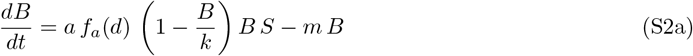

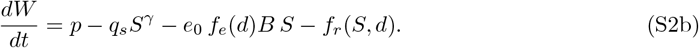

The independent and dependent variables of this dynamical system have the following dimensions:

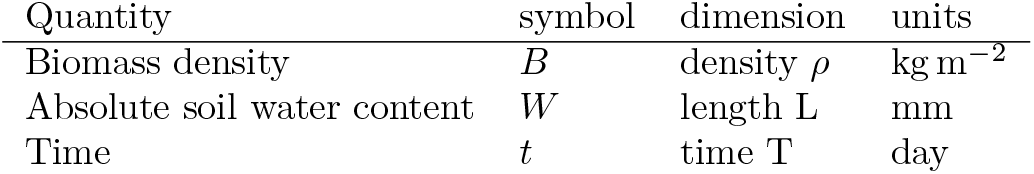

Accounting for all the model parameters that have dimensions, we have eight: *a, k, m*, (*nz*_*r*_), *p, q*_*s*_, *e*_0_, *λ*. (Here we treat *nz*_*r*_ as a single parameter.) The parameter *λ* appears inside the function *f*_*r*_(*S, d*). All of these parameters have dimensions that derive from the three basic ones shown in the table. According to Buckingham’s Π Theorem, when performing non-dimensionalization of the equations, the number of independent dimensionless groups is equal to the number of dimensional parameters minus the number of fundamental dimensions. Since we have 8 dimensional parameters and 3 fundamental dimensions (biomass density, length, and time), we obtain 8 − 3 = 5. In simple terms, although the model contains eight dimensional parameters, its behavior can be fully captured by just five independent dimensionless combinations.

We start now the non-dimensionalization process by defining the following new non-dimensional variables:

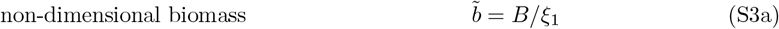

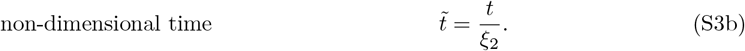

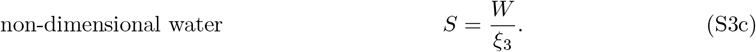

We have many choices for *ξ*_1_, *ξ*_2_, *ξ*_3_. A suitable choice here is

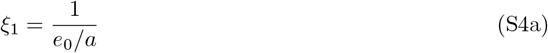

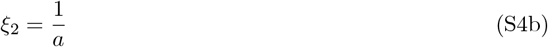

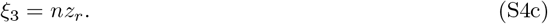

Substituting Eqs. (S3) and (S4) into Eqs. (S2) yields the non-dimensional dynamical system:

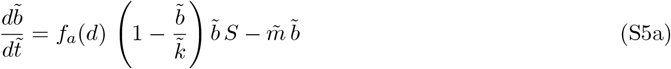

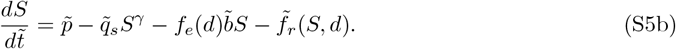

As we can see, this choice of *ξ*_1_, *ξ*_2_, *ξ*_3_ leaves us a non-dimensional system, where every single variable, parameter, term or function is dimensionless. For this specific choice of *ξ* scaling factors, we have a system with unity growth rate and unity maximum evapotranspiration rate.

The non-dimensional parameters (Π parameters) for the equations above are:

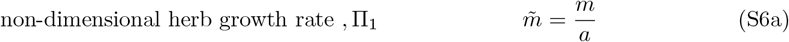

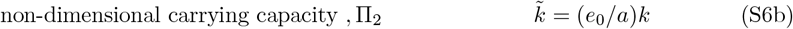

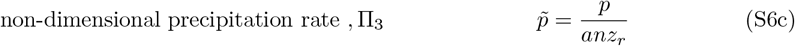

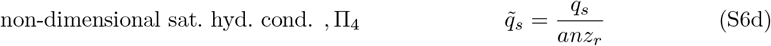

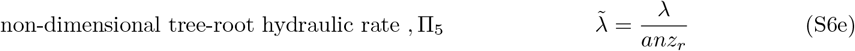

This exercise is instrumental in shedding light on the impact of various model parameters on its behavior:

- The active soil layer *nz*_*r*_ (mm), where *n* is soil porosity (dimensionless) and *z*_*r*_ is the rooting depth of herbs (mm), sets the maximum volume of water that can be stored and made available to herbs. As Eqs. (1) indicate, *nz*_*r*_ can only impact the transient dynamics of (*B, S*), never their steady-state solutions. Of course, due to the third conversion in Eq. (S3), *nz*_*r*_ rescales *S* into *W* = *nz*_*r*_*S*.
- The denominator *anz*_*r*_ represents a characteristic water throughput, where *a* sets the characteristic timescale and *nz*_*r*_ is the volume of water per unit area (1 mm = 1 L mm^*−*2^). It sets the scale for precipitation, drainage, and root uptake/uplift 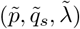: higher values correspond to processes faster than the characteristic resource throughput in the system.
- The ratio *e*_0_*/a* naturally emerges as a water-use efficiency, converting herb water uptake into biomass production. In fact, when tree density is zero (*d* = 0) and herb biomass is low (so that 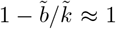), the evapotranspiration loss term in Eq. (S5b) is identical to the biomass growth term in Eq. (S5a). This symmetry highlights the direct coupling of growth and water loss. The conversion dictated by (S3a) means that whatever the non-dimensional steady state 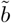 one gets by solving Eqs. (S5), we divide it by the water-use efficiency *e*_0_*/a* to get the dimensional biomass *B*.
- The typical growth timescale 1*/a* acts as a fundamental clock for all other remaining parameters (see (S6)).
- 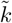shows how the dimensional carrying capacity *k* is rescaled by water-use efficiency *e*_0_*/a*. A system with a large *k* but low water-use efficiency is effectively constrained, while high water-use efficiency inflates the effective carrying capacity.

### S4 Transient dynamics

The typical timescale for the dynamics of herb biomass (*B*) is comparable to the length of the growing season used in this study, 180 days. Figure S3 shows the time dynamics (left column) and phase space (right column) for three precipitation levels (*p* = {150, 300, 600} mm year^*−*1^) and three tree conditions: no tree (*d* = 0, top row), shading mechanism (*d* = 0.3, middle row), and root mechanism (*d* = 0.3, bottom row). Linear stability analysis of the numerically found steady-state solutions (hollow circles) reveals them to be stable nodes (real and negative eigenvalues).

**Figure S3.**
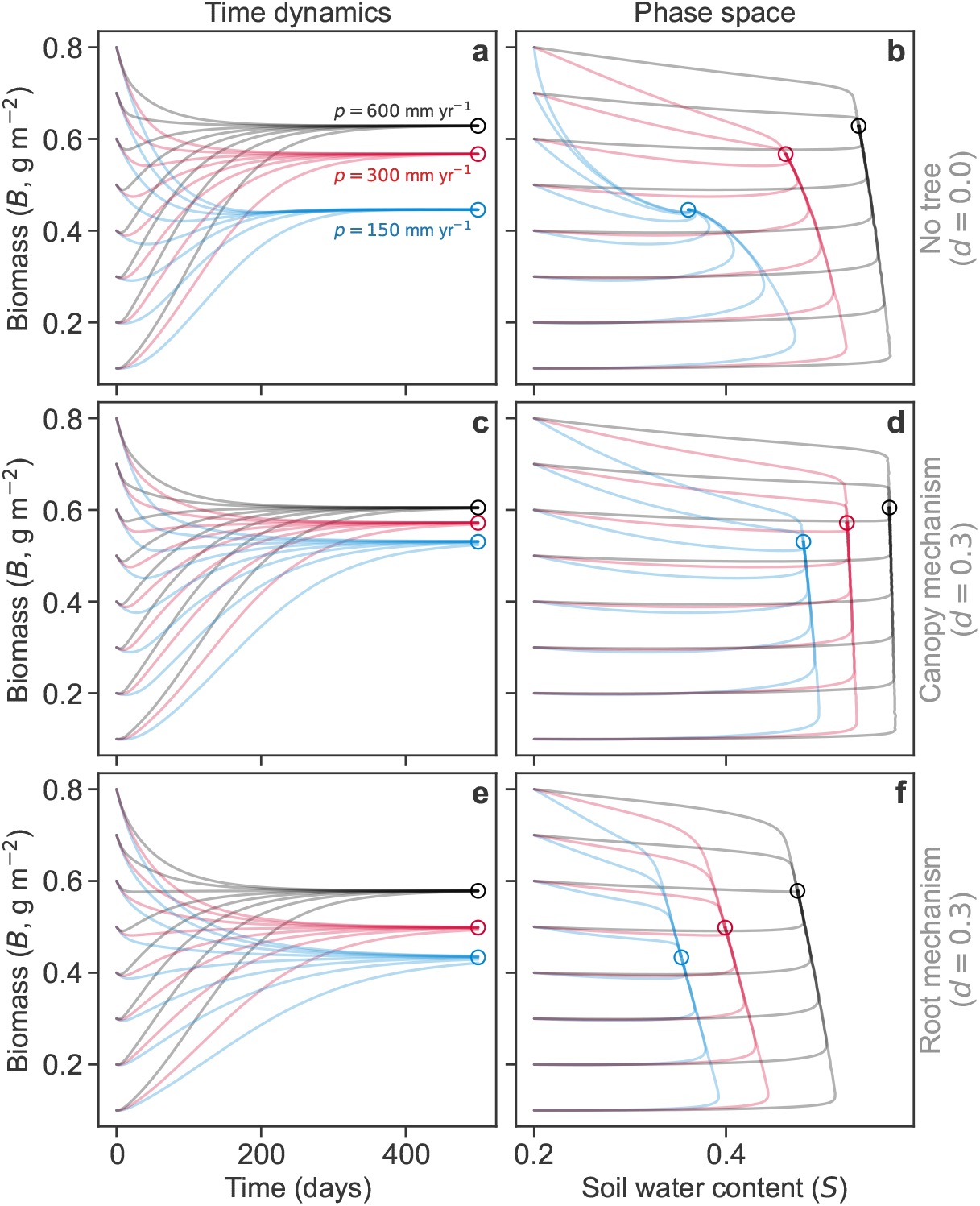
Time dynamics (left) and phase-space portrait (right) of (*B, S*) solutions. Panels a,b: zero tree density. Panels c,d: Tree density *d* = 0.3, shading mechanism on. Panels e,f: Tree density *d* = 0.3, root mechanism on. Blue, red and black orbits denote precipitation levels of 150, 300 and 600 mm year^*−*1^, respectively. The same eight initial conditions were used for all cases: *S* = 0.2 and *B* ranging from 0.1 to 0.8, with 0.1 increments. Hollow circles denote the value of the solution with initial condition (*B* = 0.4, *S* = 0.2) at time 500 days.

Crucially, the facilitation–to–competition switch persists when we examine transient dynamics, not only steady-state outcomes. In Fig. S4, we compare the steady-state pattern (panel a) with solutions obtained by integrating Eq. (1) for 180 days (panel b). The qualitative result is the same in both cases, with positive tree effects at low precipitation and negative effects at high precipitation. Temporal dynamics, however, alter the details: the magnitude of facilitation and competition, and the precipitation at which the switch occurs, differ between transient and steady-state results.

**Figure S4.**
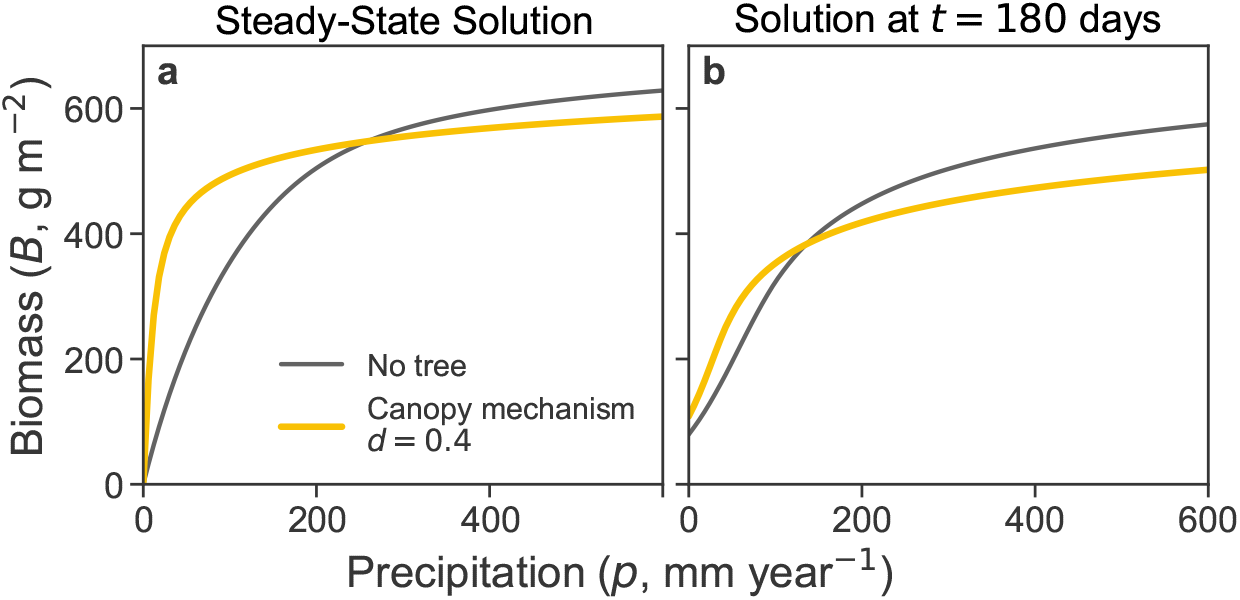
The transition from facilitation to competition, for steady-state solutions (panel a), and for transient solutions after 180 days (panel b). The initial conditions for the transient solutions are (*B* = 200 g, *S* = 0.2). For both panels we considered only the canopy mechanism, for tree density levels 0.0 and 0.4. Other parameters as reported in Table 1.

### S5 A gradient of evaporative demand

Here, we demonstrate that our model can produce a transition from facilitation to competition, not only along a precipitation gradient but also across an evaporative demand gradient (Figure S5)

**Figure S5.**
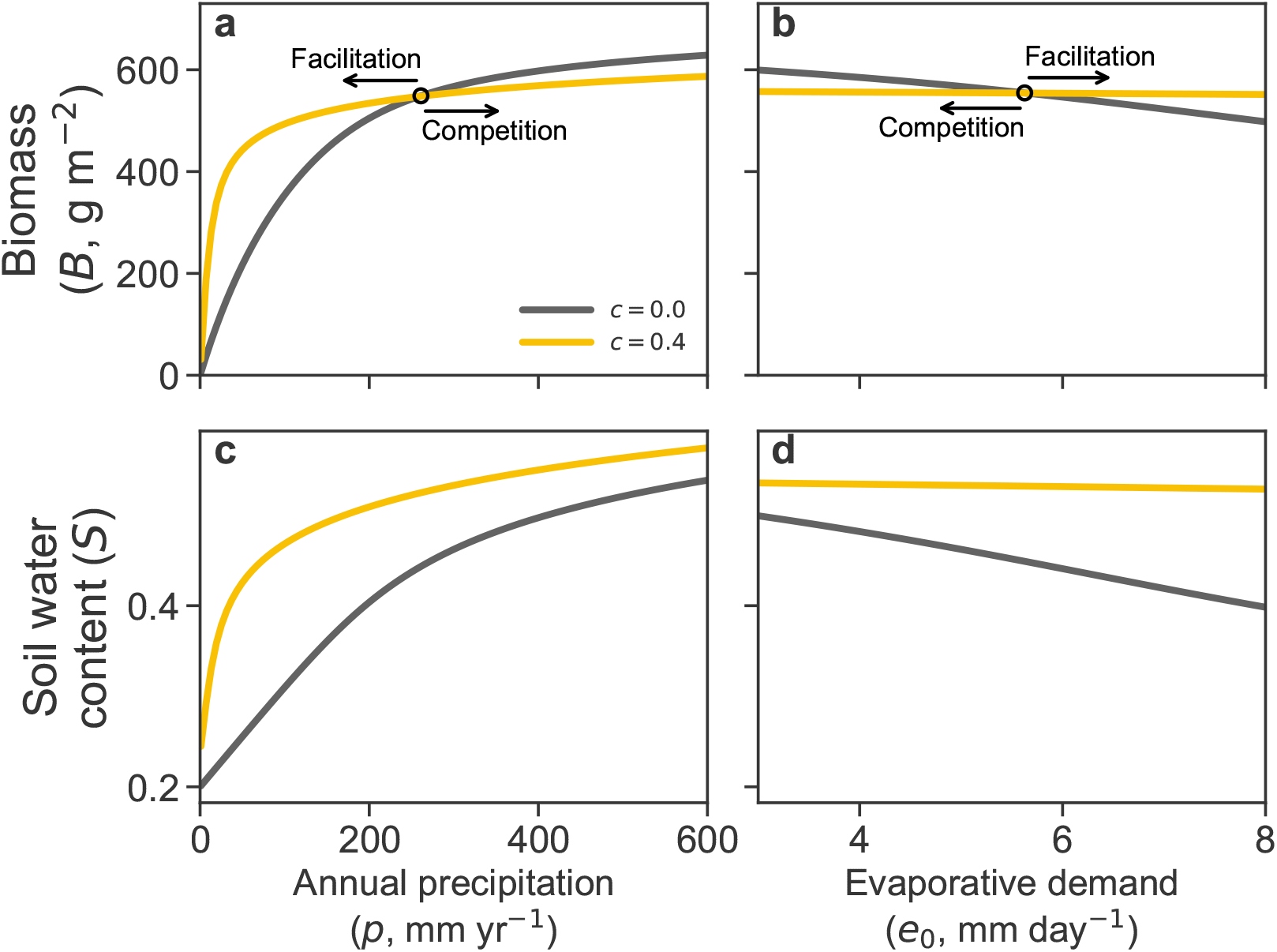
The facilitation–competition transition extends across different abiotic stress axes. Panels a,c: Precipitation as the stress axis reproduces the classic SGH prediction, with facilitation at low supply and competition at high supply. Panel b,d: Evaporative demand, representing atmospheric drivers of water loss, yields the same qualitative pattern but on a reversed axis (higher demand corresponds to stronger stress).

### S6 Choosing the Right Resource Metric for the SGH

The Stress Gradient Hypothesis (SGH) predicts a shift from facilitation to competition as a resource increases. A critical question is which resource metric to use: resource supply rate (like precipitation) or resource abundance (like soil water content). We demonstrate here that the resource supply rate is the appropriate metric for assessing the SGH.

When we analyze our model using precipitation as the *control parameter* (x-axis in a *B* vs. *p* plot), the results clearly demonstrate the SGH pattern. For both the shading and root mechanisms, we observe a transition from positive to negative tree-herb interactions as precipitation increases, mirroring field observations.

In contrast, if we use soil water content as the resource metric, the SGH pattern vanishes. Figure S6 shows the same herb biomass solutions as Figure 2, but plotted against soil water content.

**Figure S6.**
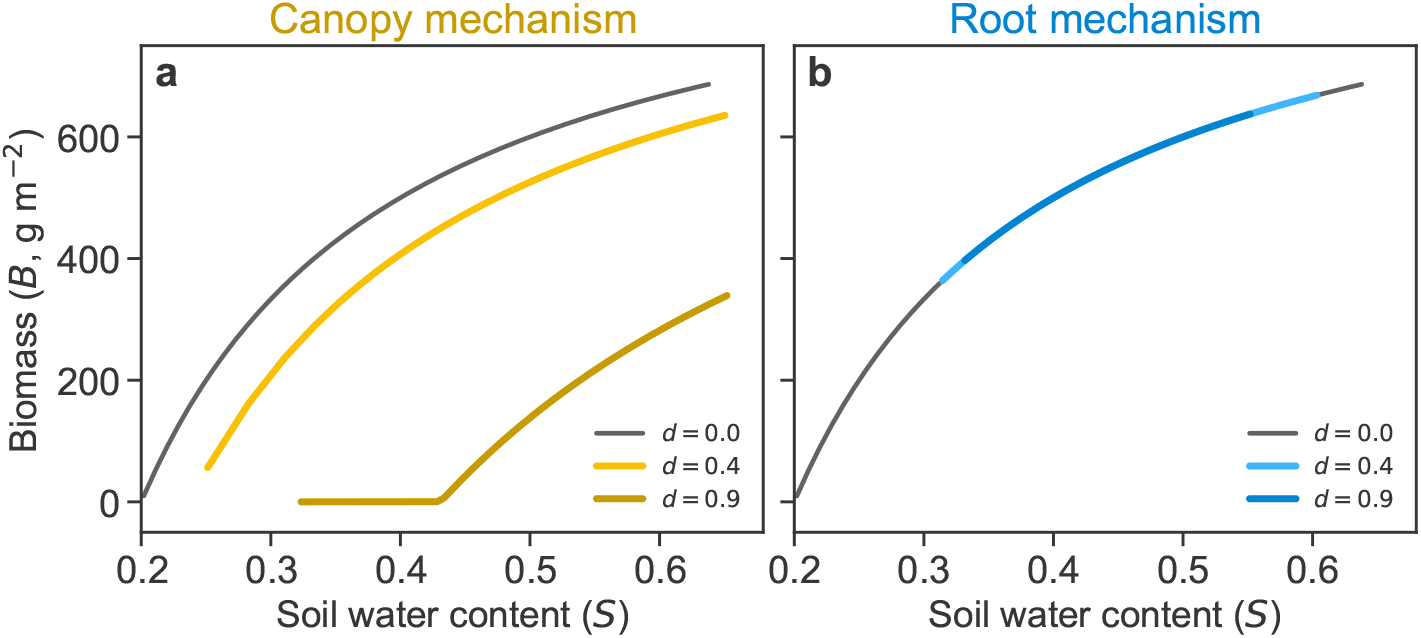
Using soil water content as the stress axis eliminates the facilitation–competition transition. Steady-state herb biomass solutions are shown as a function of soil water content for the canopy mechanism (panel a) and the root mechanism (panel b). In the canopy mechanism, biomass with trees (*d >* 0) is always lower than the no-tree baseline, because canopy directly reduces assimilation while water conservation effects are no longer represented. In the root mechanism, biomass becomes independent of tree density when soil water is fixed, since tree effects operate only by altering soil moisture levels. Accordingly, when stress is parameterized by soil water abundance rather than supply rate, the SGH pattern disappears.

The nontrivial (*B* ≠ 0) solutions can be readily obtained by equating Eq. (1a) to zero and solving for B:

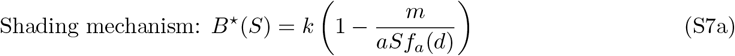

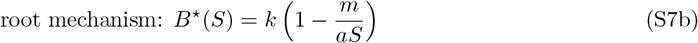

From these solutions we learn:

- **Shading mechanism:** Because *f*_*a*_(*d >* 0) *< f*_*a*_(*d* = 0), the steady-state herb biomass with trees (*B*_*T*_) is always lower than the biomass without trees (*B*_0_). Trees always reduce light, which directly suppresses herb growth. Because the facilitative effect of water conservation is no longer a factor, the interaction intensity is always negative (competitive).
- **Root mechanism:** Herb biomass becomes completely independent of tree density (it lacks the variable *d*). Trees influence herbs solely by altering soil water levels, so if that level is fixed, the trees have no effect. The interaction intensity is always zero.

This result, that the SGH pattern disappears when using soil water content as the metric, occurs for a fundamental reason: soil water content, unlike precipitation, is an intrinsic property of the ecosystem, not an external force. It is the result of multiple processes such as rainfall, evapotranspiration, drainage, etc; all of which interact with the tree’s presence to determine the final soil water content. By holding soil water constant, we are artificially decoupling the very mechanisms that create the SGH pattern. Therefore, using resource abundance (soil water content) as the metric of stress renders the SGH meaningless within this mechanistic framework, as it nullifies the resource component in our **consumer-resource model**.

The **resource supply rate** (precipitation) is the external driver of the system’s state, making it the correct measure of abiotic stress for a resource-based SGH. It allows us to capture the full interplay of facilitative and competitive forces that are the core of the hypothesis.

### S7 Logistic growth extension: nutrient-limited growth

To justify our choice of introducing a limiting factor to herb growth as a logistic term, we do the exercise of explicitly introducing a new essential resource, a nutrient *N*. Our model can be extended to include the nutrient dynamics as follows:

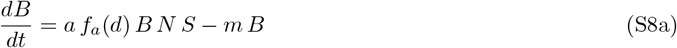

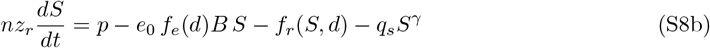

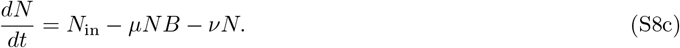

The first equation, (S8a), describes the herb biomass dynamics, where we have replaced our original logistic term *f*_*k*_(*B*) with a nutrient term, *N*. Equation (S8c) describes the nutrient’s dynamics, where it is introduced at a constant rate *N*_in_, consumed by herbs (at a rate proportional to both herb biomass and nutrient availability, *µNB*), and lost from the system at a rate proportional to its own abundance, *νN*.

We now assume that the nutrient dynamics are much faster than those of the herb biomass (*B*) and soil water (*S*). We can therefore perform an adiabatic elimination by setting the rate of change of the nutrient to zero to find its quasi-steady-state:

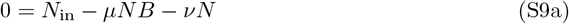

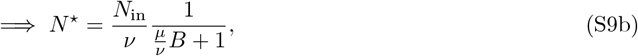

where *N* ^*⋆*^ denotes the quasi-steady-state nutrient concentration. Substituting *N* ^*⋆*^ into the herb biomass equation (S8a) gives:

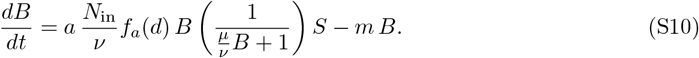

The term in the parenthesis acts as a saturating function of *B*. When herb biomass is low, the term is close to one, and as biomass increases, the function decreases, effectively slowing the growth rate. This plays the same role as the logistic term we assumed in the main text.

To derive the exact logistic form, we can make an additional assumption. If we assume that the herbs are highly inefficient at consuming the nutrient relative to its decay rate (*µ* ≪ *ν*), we can use a first-order Taylor series expansion:

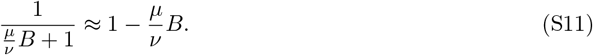

Substituting this approximation into the herb biomass equation now reveals a classic logistic term:

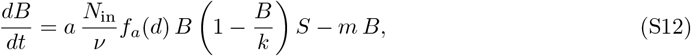

where the carrying capacity can be identified with *k* = *ν/µ*. If the factor *N*_in_*/ν* is then incorporated into *a*, we get exactly Equation (1a) in the main text.

### S8 Tree density dependent on precipitation

The arguments in Supplementary Section S1 stated that there can be no SGH transition in the absence of a carrying capacity term for constant tree density. This picture changes if instead we allow for precipitation-dependent tree density (as observed in many systems). Higher precipitation supports greater tree density, which, in turn, exerts a stronger shading impact on herbaceous species. We consider here two distinct functional forms of *d*(*p*):

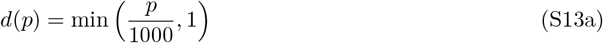

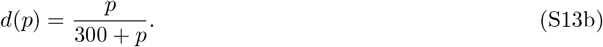

While the model internally uses *p* in mm d^*−*1^, we present the equations *d*(*p*) in mm y^*−*1^ for simplicity and practical relevance.

Figure S7a illustrates the transition from facilitation to competition for these two functional forms. Figure S7b shows the absolute interaction intensity in the (*p, d*) plane, and helps us interpret the graph on the top. When a dark curve in the bottom panel crosses the border between facilitation and competition, we see in the top panel that the dark curves cross the no-tree solution. For higher values of *d*, when the dark curves in the bottom panel cross into the no-viable-biomass zone, we also see a collapse of the vegetated solutions in the top panel. Finally, Fig. S7b helps us understand, from another point of view, why there cannot be a transition from facilitation to competition in the case of no carrying capacity and constant tree density, as previously discussed. Constant density solutions (see for example the dotted line at *d* = 0.2) always cross from competition to facilitation as *p* increases, never the other way around. By changing tree density from a constant value to increasing functions of precipitation, as depicted by the solid dark lines, one can get the usual transition pattern of the SGH.

**Figure S7.**
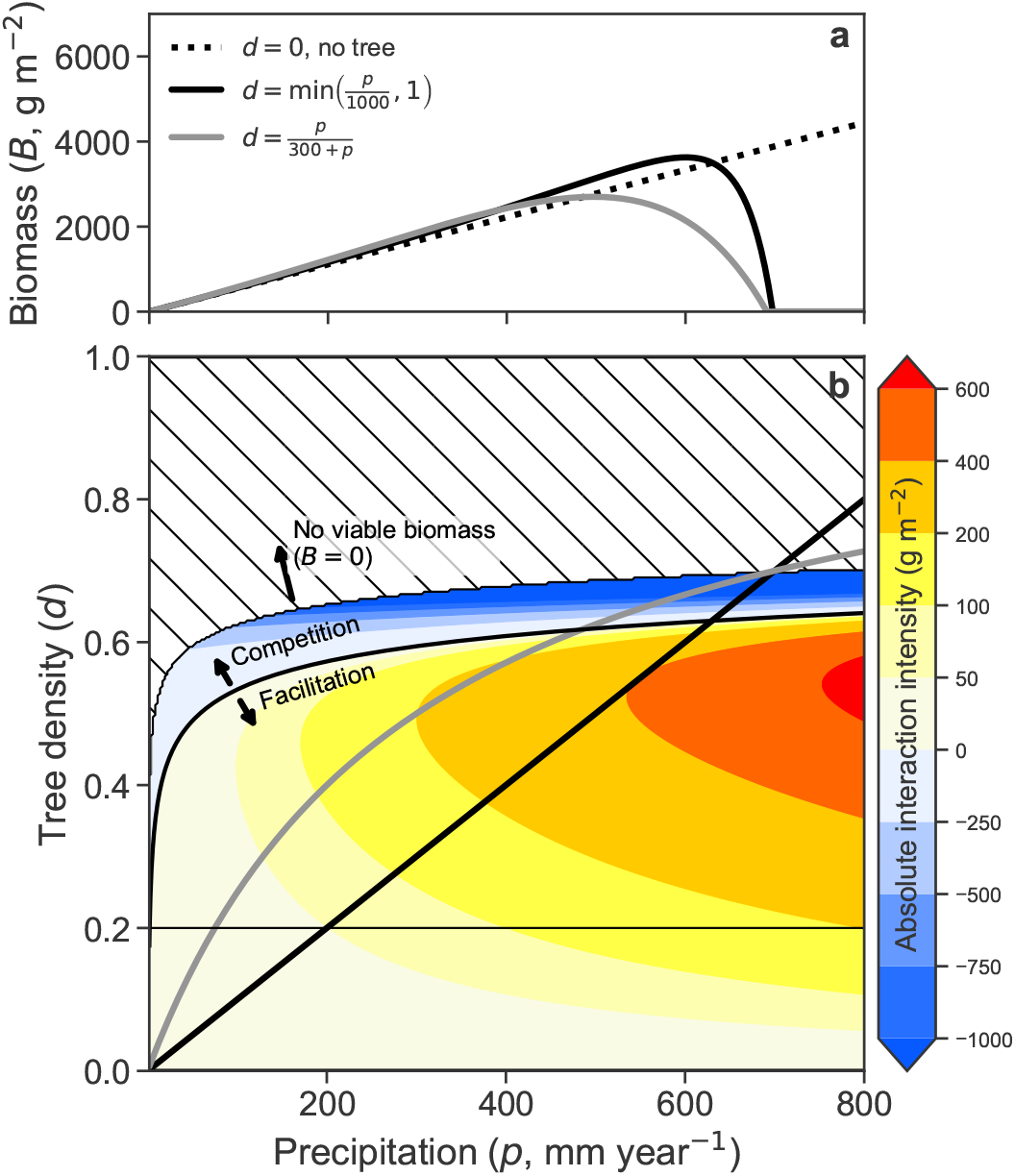
The Precipitation-Dependent Tree density Mechanism produces the SGH transition even without a carrying capacity term. Panel a: Steady-state herb biomass solutions (Eq. (S1b)) for a scenario without any tree density (dotted line) and for two possible instantiations of the tree-density dependence on precipitation (dark curves). Panel b: The absolute interaction intensity is shown in the (*p, d*) plane: warm colors denote positive interactions (facilitation), while cold colors denote negative interactions (competition). The hashed region in the top indicates no viable herb solutions (*B* = 0). The two functional forms for tree-density are plotted in solid dark curves. Parameters: *β*_*a*_ = 0.9 and *β*_*e*_ = 1.1, and other parameters as shown in Table 1.

### S9 Infiltration

The infiltration mechanism is a variation of the canopy mechanism. In both, canopy shading reduces herb assimilation through *f*_*a*_(*d*); here, however, the facilitative pathway is not a reduction in evapotranspiration but an increase in effective infiltration as tree density rises. The model therefore keeps the growth-side shading term and replaces the rainfall input with an infiltrated input *p f*_*i*_(*d*):

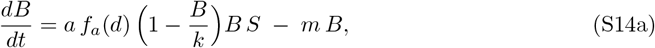

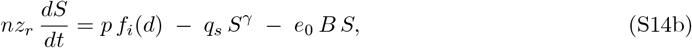

The infiltration factor *f*_*i*_(*d*) follows ^2^:

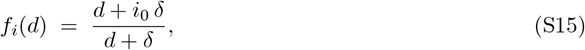

so that *f*_*i*_(0) = *i*_0_ (minimum infiltration without trees) and *f*_*i*_(*d*) *→* 1 as *d* increases; see Fig. S8a. The parameter *δ* sets the density scale at which *f*_*i*_ is midway between *i*_0_ and 1. Figure S8b shows the SGH transition for the infiltration mechanism.

**Figure S8.**
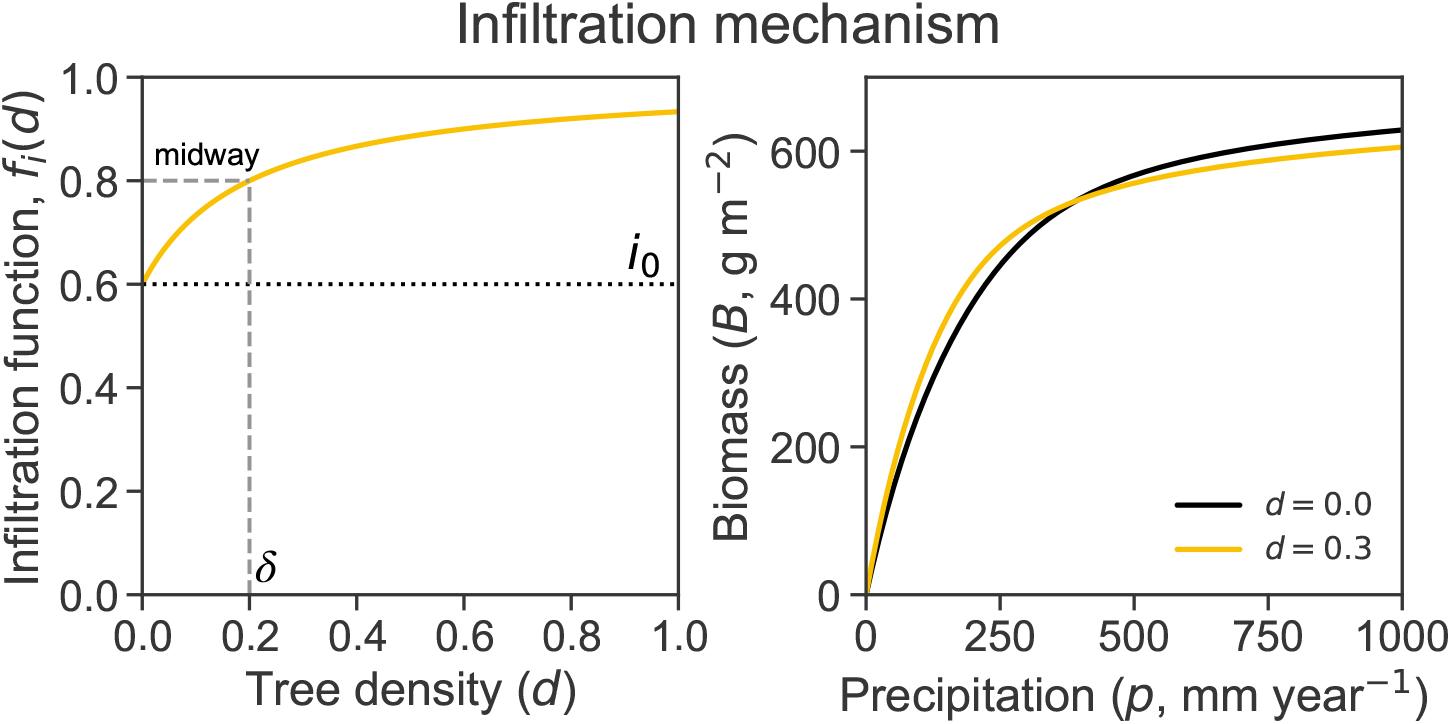
Infiltration mechanism as a canopy variant. (a) The infiltration factor *f*_*i*_(*d*) increases from *i*_0_ at *d* = 0 toward 1 as tree density rises (*δ* controls the transition scale). (b) The utransition from facilitation to competition can be obtained from the infiltration mechanism.

This relationship captures the idea that canopy shading can suppress biological soil crusts (biocrusts), which often form a hardened surface layer that reduces infiltration. As tree density increases, shading limits biocrust activity and allows more water to infiltrate into the soil, thereby increasing the effective water supply to herbs.

## Footnotes

2 Slope σ: for d = 0 the slope is σ_0_ = a/(e_0_m), whereas for d > 0 it is σ_d_ = σ_0_ · (f_a_(d)/f_e_(d)). Since f_a_(d) > f_e_(d), we have that σ_d_ > σ_0_. Intercept ω: for d = 0 the intercept is ω_0_ = −(q_s_/e_d_)(m/a)^γ−1^, whereas for d > 0 it is 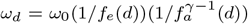. Since f_a_(d), f_e_(d) < 0 and γ > 2, we have that each of the parenthesis in the expression for ω_d_ is greater than 1, and therefore ω_d_ < ω_0_ (remember the minus sign in ω_0_).

